# Visualising sub-second dynamics of nanoparticle extravasation *in vivo*

**DOI:** 10.1101/2025.08.10.669578

**Authors:** Guoying Wang, Yueying Cao, Jia Li, Xianlin Zheng, Peng Ren, Marco Morsch, Bingyang Shi, Yiqing Lu

## Abstract

Altered vascular integrity is a hallmark for many diseases. Monitoring vascular permeability, including its extent and underlying transport pathways, is thus important for advancing our understanding of disease mechanisms towards innovative diagnosis and treatment. Here we show real-time fluorescence imaging using lanthanide-based upconversion nanoparticles as contrast agents to visualise subtle changes to vascular permeability *in vivo*. Based on simultaneous confocal imaging of the vasculature alongside single particle tracking in the wide-field at video rate, we performed high-throughput surveillance across live zebrafish larvae to pinpoint locations of potential nanoparticle extravasation, achieving superior sensitivity and specificity over conventional fluorescent dyes. Further analyses of the sub-second dynamics of individual nanoparticle extravasation events unveiled distinct characteristics to distinguish between transcellular and paracellular transport. We applied the technique to evaluate the blood–brain barrier (BBB) in zebrafish larvae at different developmental stages, and potential BBB perturbation strategy via nanoparticle functionalisation with polysorbates. Significantly increased BBB penetration by 36.5 folds was shown for the functionalised particles compared to phospholipid-coated particles, attributed to enhanced transcellular crossing. The technique is readily applicable to monitoring vascular integrity and investigating endothelial transport for improved understanding of vascular biology, facilitating advanced research in disease diagnostics and drug delivery towards translation.

## Introduction

Investigation into vascular permeability is attracting increasing attention due to its importance in physiology, pathology, diagnostics and therapy^1-4^. In Alzheimer’s disease for instance, disrupted integrity of the blood–brain barrier (BBB) is one of the earliest markers and contributes to subsequent cognitive decline^2,5,6^. Stroke and cerebral cavernous malformation also result in BBB breakdown^7,8^, whereas repairing BBB integrity is regarded as a promising therapeutic approach for various brain disorders^9,10^. Moreover, distinguishing the underlying pathways between transcellular and paracellular transport is deemed crucial in addition to the extent of permeability, as emerging evidence indicates these mechanisms play distinct roles in the progression of brain pathology such as stroke and aging^7,11^. The capability to differentiate transport pathways is also in urgent need to improve targeted drug delivery such as cancer nanomedicine and emerging pharmaceuticals based on extracellular vesicles^12,13^. These underscore the critical importance of monitoring vascular permeability and elucidating the precise pathways involved in order to better understand disease mechanisms and drive the development of innovative diagnosis and treatment.

Such targeted observations, however, have been hampered by the lack of suitable methodologies. Many studies have relied on *ex vivo* examination, which retains no dynamic information^14^ and can potentially introduce artefacts during sample preparation. Alternatively, medical imaging has been used to investigate vascular permeability *in vivo* (i.e. angiography)^15,16^ but can only reveal relatively severe, most likely paracellular leaks^16,17^. By contrast, real-time imaging with fluorescent probes provides a simple and accessible approach, offering good spatial and temporal resolution^3,17,18^. Although the sensitivity of conventional fluorophores has previously been an issue, this may be addressed by using nanoparticle probes as contrast agents to facilitate single particle tracking (SPT)^19-30^. Nevertheless, a fundamental trade-off remains between temporal and spatial performance, as increase in the frame rate is typically at the expense of the number of pixels (thereby reducing either the field-of-view or the spatial resolution), especially for laser-scanning confocal and multi-photon microscopy where adequate pixel dwell time has to be maintained to assure single nanoparticle sensitivity. The effective frame rate of SPT further decreases in the case of optical sectioning to acquire vasculature information in three dimensions. Alternatively, wide-field fluorescence microscopy can provide continuous observation of single nanoparticles, but appeared to be applied mainly *in vitro* to date, presumably due to considerable interference from biological autofluorescence in thick specimens. New methods remain to be developed to unveil minute changes to the vascular permeability along with the exact microscopic pathways *in vivo*.

We propose that, the key to unlocking the desirable capability lies in the disentanglement of fast nanoparticle imaging required for SPT from vasculature imaging potentially at lower speed, while still performing both concurrently to provide real-time environmental context of the nanoparticle movement. Specifically, we employed lanthanide-based upconversion nanoparticles (UCNPs), whose anti-Stokes signal is readily separable from Stokes-shifted normal fluorescence, and developed a correlative wide-field (for particle tracking) and confocal (for vasculature imaging) microscopy platform (see Methods). This technique allowed us to capture the real-time trajectories of single nanoparticle extravasation in live zebrafish larvae, and analyse their dynamic features to distinguish between transcellular and paracellular transport. We demonstrated the advantages of using nanoparticle probes over conventional fluorescent dyes in terms of sensitivity and specificity. We further applied the technique to evaluate the BBB integrity in zebrafish at different developmental stages, and the influence of nanoparticle surface functionalisation on BBB penetration.

## Results

To visualise vascular permeability, we injected UCNPs into a transgenic zebrafish model [Tg(fli1a:EGFP)]^31,32^ and imaged the sample on the purpose-built microscopy platform (Fig. 1a). When continuously imaged in the wide-field, these nanoparticles exhibited exceptionally photostable photoluminescence, with their brightness optimised^33-35^ to achieve single particle sensitivity at 20 full frames per second (Fig. 1b). Since the upconversion emission from UCNPs was easily separated from the fluorescence signal of common fluorophores, vasculature imaging can be performed simultaneously for the same field-of-view to provide the spatial context of the nanoparticle movement (Fig. 1c). This dual-modal imaging technique enabled high-throughput surveillance of one fish sample in 2 minutes to quickly identify and localise permeable blood vessels, if any, followed by high-speed recording of the nanoparticle extravasation process (Fig. 1d). Co-registration of the UCNPs with respect to the vasculature at high resolution in 3D further allowed quantitative analysis of the nanoparticle biodistribution *in vivo*, offering insights into the extravasation probability at specific sites across the living organism (Fig. 1e).

**Figure 1.**
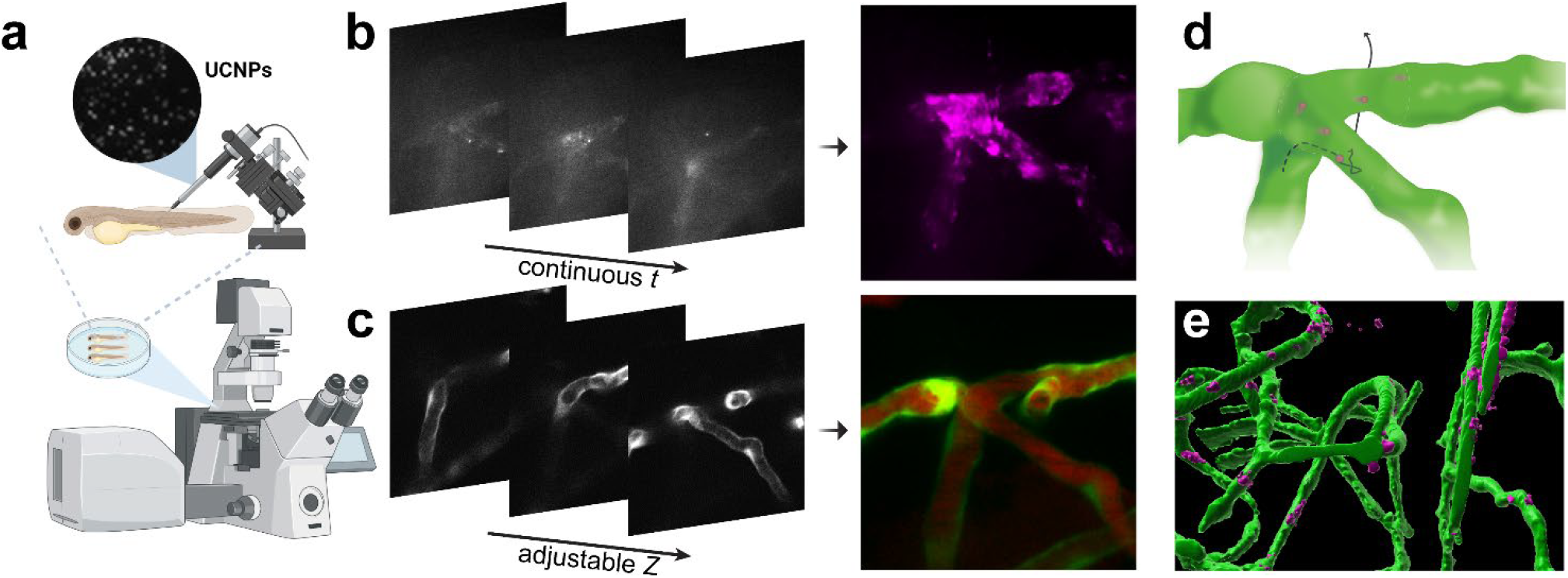
Real-time fluorescence imaging using UCNPs as contrast agents to assess vascular permeability in live zebrafish. (a) Zebrafish injected with UCNPs in the vasculature are imaged on a purpose-built microscopy platform. (b, c) Continuous wide-field imaging to track UCNPs (rendered in magenta) over time, and simultaneous confocal imaging of the zebrafish vasculature (rendered in green, whereas the red signal is from a fluorescence dye to indicate the lumen) on the same focal plane. The focus is adjustable in real time with Z position continuously recorded during the dual-modal imaging. (d) Construction of the dynamic trajectory of a single nanoparticle with respect to the blood vessels. (e) 3D co-registration of the UCNPs and the blood vessels.

Upon successful validation of single nanoparticle sensitivity *in vitro* (see Supplementary Notes 1 & 2), we implemented the technique to distinguish the *in vivo* pathway of nanoparticle extravasation based on the real-time dynamics. In the brain area of 3 dpf (days post fertilisation) old zebrafish larvae (at which stage endothelial tight junctions are functional, but transcytosis is not fully regulated^36^), we observed UCNP extravasation across the BBB in a typical pattern of slow and confined movement before exiting the vasculature (Fig. 2a and Supplementary Movie 1). In comparison, UCNP extravasation from the intersegmental vessel in the zebrafish trunk generally manifested itself as rapid passage followed by Brownian motion outside the vessel (Fig. 2b and Supplementary Movie 2). Quantitative analyses of the trajectories revealed several distinctive parameters corresponding to potential transcytosis vs paracellular crossing. In the former (Fig. 2c-e), the nanoparticle co-localised with the vessel wall for an extended duration as long as 10 minutes since being detected, during which period it exhibited slow (< 1 μm/s) but directed motion (Phase 1) from time to time. This behaviour was likely attributed to intracellular movement along structures such as the actin filament, suggesting nanoparticle uptake by the endothelial cell. The displacement then started to increase as the nanoparticle traversed the BBB and entered the brain parenchyma (Phase 2), often orientated towards certain direction, with the instantaneous speed remaining relatively low (< 2.8 μm/s). The two phases were also marked by their mean squared displacement (MSD), indicating Phase 1 was sub-diffusive (MSD exponent < 1) and Phase 2 was directed (MSD exponent > 1). In the latter (Fig. 2f-h), the nanoparticle displayed significantly higher peak speed (> 5 μm/s), while the direction of movement was more random with an MSD exponent close to 1, characteristic of Brownian motion. These dynamic features provide unprecedented evidence *in vivo* to elucidate the transport pathways distinguishing between transcellular and paracellular extravasation.

**Figure 2.**
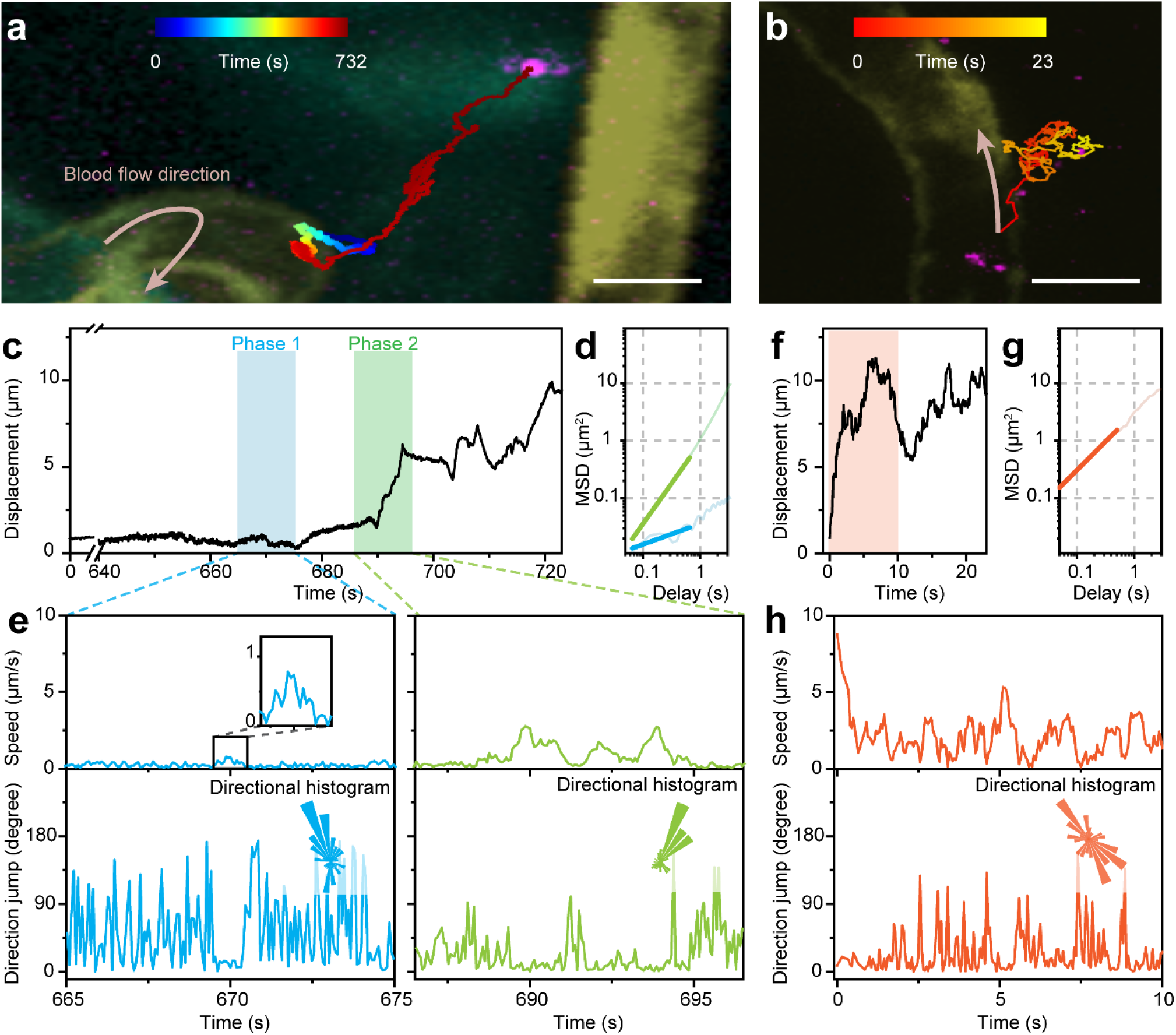
Distinguishing between transcellular and paracellular extravasation based on the dynamic trajectories of individual nanoparticles. (a) A representative extravasation event in the brain of 3 dpf zebrafish, corresponding to Supplementary Movie 1. The trajectory is colour-coded in time and superimposed on the confocal image that is colour-coded in depth. (b) A representative extravasation event in the trunk of 3 dpf zebrafish, corresponding to Supplementary Movie 2. (c) Displacement of the nanoparticle in (a) over time. The two phases before and after extravasation are shaded in blue and green, respectively. (d) MSD plot in the log-log scale for the two phases, with fitted slopes (i.e. MSD exponent) of 0.4 for Phase 1 and 1.4 for Phase 2. (e) The instantaneous speed and the direction jump over the enlarged periods in (c). Insets in the direction jump sub-plots are the directional histograms of the particle movement in the corresponding phases. (f) Displacement of the nanoparticle in (b) over time. (g) MSD plot in the log-log scale over the orange-shade period in (f), with a fitted slope (i.e. MSD exponent) of 1.0. (h) The instantaneous speed and the direction jump over the orange-shade period in (f), with the inset being the directional histogram. Scale bars: 10 μm.

To demonstrate the potential advantages of using nanoparticle contrast agents in examining vascular permeability, we injected both UCNPs and dextran-tetramethylrhodamine isothiocyanate (dextran-TRITC, 10 kDa) in the zebrafish vasculature to evaluate the BBB integrity for comparison. In multiple samples of 3 dpf zebrafish larvae, UCNPs were observed outside the BBB, with the extravasation sites clearly located by the recorded trajectories (Fig. 3a). The signal-to-background ratio for individual UCNP (derived from the peak intensity in the white box containing the particle divided by the mean value across the field-of-view) exceeded 5.2 in each frame (Fig. 3b), allowing unambiguous identification and continuous tracking of the nanoparticle. By contrast, the fluorescence signal of dextran-TRITC appeared insignificant at the same spot, with signal-to-background ratio of 1.4 that lacks the sensitivity to locate the site of compromised BBB integrity (Fig. 3c). In addition, higher-than-average background was observed in other areas (e.g. the blue box) for the dextran-TRITC channel, suggesting inferior specificity using conventional fluorescent probes compared to the nanoparticles. In 5 dpf zebrafish, we observed no nanoparticle extravasation events (Fig. 3d), but a considerable amount of dextran-TRITC distributed in the brain parenchyma (Fig. 3e). A thread-like pattern was seen connecting the brain parenchyma to the bright region around the front head surface vessels, suggesting that instead of BBB leakage, the dextran-TRITC was extravasated through vessels outside the BBB^37^. We performed more control experiments using another widely applied fluorescent probe for BBB integrity, Alexa Fluor 647 labelled BSA (BSA-AF647, 66 kDa). A different pattern was observed, showing aggregates accumulated both along and outside the vasculature (Fig. 3f). Nonetheless, some BSA-AF647 was also observed around the front head surface vessels, so that it remained inconclusive whether the probe entered the brain via BBB crossing or not. The comparative results confirmed that tracking nanoparticles offers higher sensitivity and specificity to precisely determine the location, timing, and pathway of potential BBB penetration events, which is not feasible for conventional fluorescent probes when used solely.

**Figure 3.**
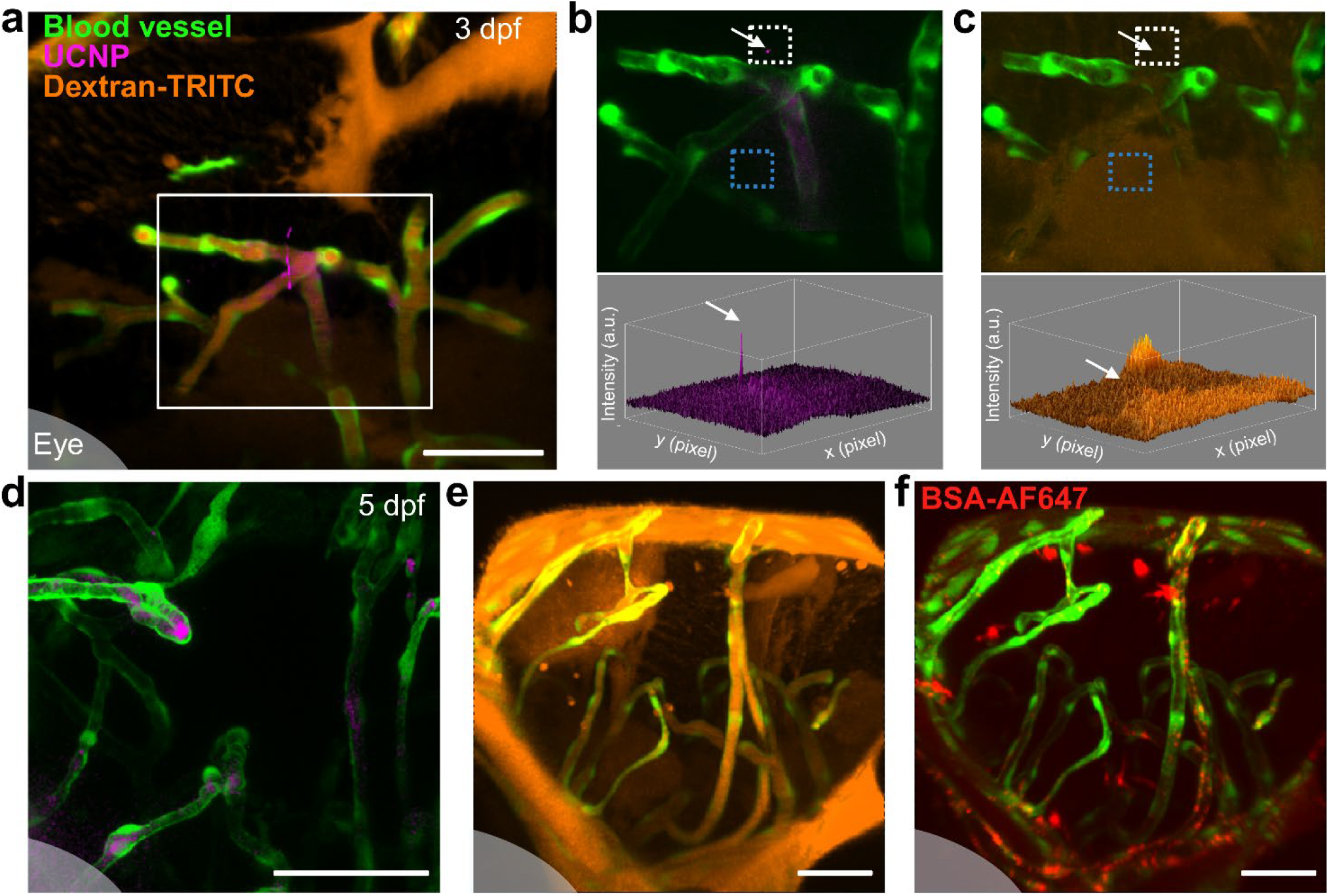
Comparison between UCNPs and fluorescent dyes for the assessment of BBB integrity. (a) A representative image in the brain of 3 dpf zebrafish showing UCNPs (magenta) and dextran-TRITC (orange) in relation to blood vessel (green). (b) The UCNP channel of the area outlined in (a) and the corresponding intensity profile. (c) The dextran-TRITC channel of the same area and the corresponding intensity profile. The white arrows indicate the same position corresponding to the extravasated UCNP. The areas marked by white and blue dotted strokes are used for calculating the signal-to-background ratio. (d-f) Representative images of the brain in 5 dpf zebrafish, captured 1 h post-injection of the UCNPs, dextran-TRITC, and BSA-AF647 (red) as contrast agents. The shaded area at the bottom left corner in (a & d–f) indicates the zebrafish eye. All scale bars: 50 µm.

We applied the technique to further explore the progressive changes in BBB integrity against various factors. For example, in 5 dpf zebrafish, no BBB penetration event was ever observed over 23 fish samples (except for the area around circumventricular organs between mid- and hindbrain), indicating a matured BBB at this developmental stage (Fig. 4a). Conversely, BBB permeability was prominent in 18 fish samples at 3 dpf (Fig. 4b) and all the captured nanoparticles were recognised as having transcellular-type dynamics (Fig. 4c-d), which is consistent with incomplete transcytosis regulation at this stage^36^. The absence of paracellular transport events being observed supports the consensus that the tight junctions among brain endothelial cells stop nanoparticles from passing the BBB in general, whereas deliberate damaging the BBB via laser ablation resulted in a large number of nanoparticles exemplifying paracellular extravasation. We then evaluated how nanoparticle surface functionalisation could affect the BBB penetration, in particular with polysorbate 80 (PS80), which was known to induce BBB passage but the mechanism was a matter for debate^18,38^. Indeed, we found that the PS80-functionalised UCNPs showed much higher probability to pass the BBB than the original phospholipid-coated UCNPs in both 5 dpf (Fig. 4e) and 3 dpf (Fig. 4f) zebrafish. Moreover, all the observed nanoparticles demonstrated dynamic trajectories corresponding to the trans-endothelial route (Fig. 4g-h), and more PS80-functionalised UCNPs were found colocalised with the vessel walls (Fig. 4i) compared to phospholipid-coated UCNPs (Fig. 4j). Quantitative analyses of the UCNP intensity showed 21-fold higher overall retention in the brain region and about twice the percentage of particles (80% vs 46%) entering the brain parenchyma (Fig. 4k), yielding a 36.5-fold increase in the quantity of particles crossing the BBB. These data strongly support that the primary contribution of PS80 arises from increased transcytosis via the brain endothelial cells rather than disrupting the BBB tight junctions^39^.

**Figure 4.**
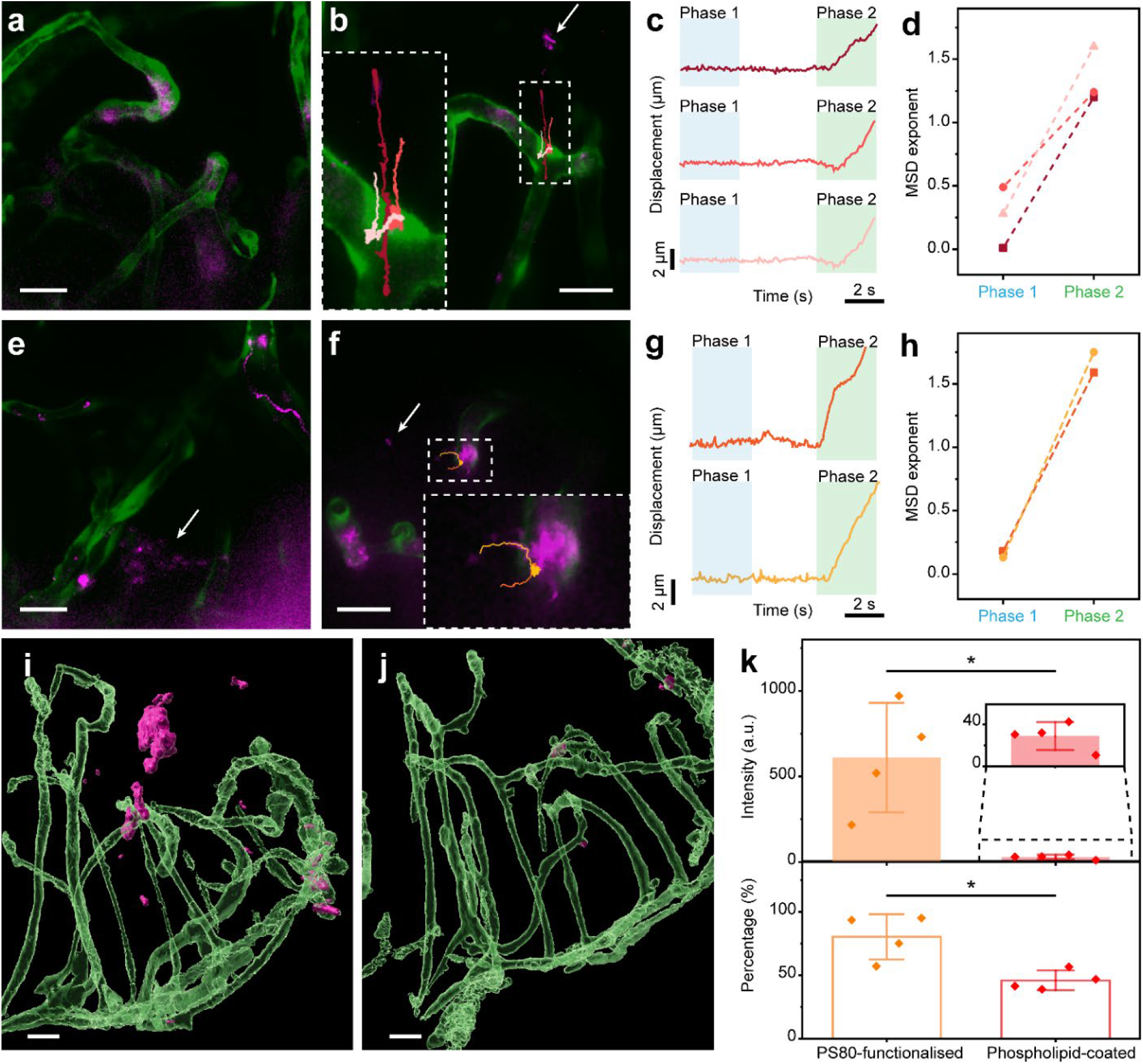
BBB permeability in relation to brain developmental stage and nanoparticle surface functionalisation. (a, b) Co-registration of the confocal and the wide-field images in the brain of 5 dpf and 3 dpf zebrafish, respectively, injected with phospholipid-coated UCNPs. The arrow points to UCNPs in the brain parenchyma, and the recorded trajectories crossing the BBB (marked in different colours) are enlarged. (c) Displacement curves extracted from the trajectories in (b), showing Phases 1 and 2 in blue and green shades, respectively. (d) The power-law-fitted MSD exponents for Phases 1 and 2 of the three displacement curves in (c). (e, f) Co-registration of the confocal and the wide-field images in the brain of 5 dpf and 3 dpf zebrafish, respectively, injected with PS80-functionalised UCNPs. The arrows point to UCNPs in the brain parenchyma, and the recorded trajectories crossing the BBB are enlarged. (g) Displacement curves extracted from the trajectories in (f), showing Phases 1 and 2 in blue and green shades, respectively. (h) MSD exponents for Phases 1 and 2 of the two displacement curves in (g). (i, j) 3D-rendered brain regions of 5 dpf zebrafish injected with PS80-functionalised and phospholipid-coated UCNPs, respectively. The blood vessels are rendered semi-transparent to help visualise the UCNP distribution. (k) UCNP intensity (total in the brain region of 1.8×10^7^ μm^3^) and percentage (outside brain vasculature to total in the region) in the brain of 5 dpf zebrafish, comparing the two groups injected with PS80-functionalised and phospholipid-coated UCNPs (*n* = 4 fish per group). Data are represented as mean ± standard deviation, with the asterisk representing *p* < 0.05 for Mann-Whitney test. All scale bars: 20 μm.

## Discussion

Combining dual-modal fluorescence imaging with single particle tracking, the technique developed in this work enables direct visualisation of altered vascular permeability and underlying transport pathways *in vivo*. The imaging platform provides continuous monitoring of nanoparticle contrast agents at 20 frames per second, facilitating rapid surveillance over an entire zebrafish larva (3–4 mm body length) within 2 mins to identify and localise permeable blood vessels. The dynamic characteristics (speed, direction jump, and MSD) obtained from individual nanoparticle extravasation events have revealed two distinct transport behaviours. The first type of events features confined motion before extravasation at relatively low speed (Supplementary Figure 1), corresponding to intracellular movement that is essential for transcytosis; the second type of events exits the circulation without intracellular retention and exhibits subsequent Brownian motion (Supplementary Figure 2), indicative of paracellular transport. Such differentiation relies on single particle tracking at high frame rate *in vivo*, which is challenging for conventional dyes. We have implemented this technique to quantify the effect of PS80 functionalisation on nanoparticle BBB penetration in zebrafish, showing 36.5-fold enhancement that is primarily through transcytosis. The visualisation and quantification capabilities demonstrated here open a new avenue to evaluating hypotheses and reconciling controversies related to vascular biology.

To enable integration of fast nanoparticle tracking into concurrent vasculature imaging, the choice of UCNP here is considered advantageous given minimum interference between the upconversion signal and normal fluorescence (including biological autofluorescence) signals. It also benefits from exceptional photostability for longitudinal monitoring^40,41^, compatibility with super-resolution imaging^42^, as well as satisfactory biocompatibility with living organisms^43,44^ and near-infrared wavelengths to assist in deep-tissue penetration^45,46^. The superior signal-to-background ratio over the course of tracking is key to improved sensitivity and specificity, avoiding potential misinterpretation when attempting to pinpoint minor leakage. The development of the dual-modal imaging platform has been made relatively straightforward by utilising a two-deck microscopic frame (Olympus IX83), with each deck configured for either confocal or wide-field imaging, as well as simplified calibration and alignment. The platform supports further integration with other fluorescence microscopic modalities (e.g. fluorescence lifetime imaging, total internal reflection fluorescence, fluorescence correlation spectroscopy) and functionalities (microinjection, laser ablation, electroporation, etc.), rendering it highly versatile to address sophisticated experimental requirements.

It should be noted that capturing the entire process of nanoparticle extravasation is not free of challenges due to the unpredictable nature of the exact location and timing. An important factor for consideration is the quantity of nanoparticles injected, which should balance the occurrence of extravasation events (especially at places with inherently low probability, such as the BBB) and the interference from other particles in the circulation. Once mastered, the new technique is readily applicable to monitoring vascular integrity and endothelial transport for improved understanding of diseases. For example, using a zebrafish model of Alzheimer’s Disease^47^, potential alteration to BBB permeability can be correlated to disease onset and progression as well as the effect of emerging therapeutic interventions aiming to restore BBB integrity, generating new information to advance pathological and bioinformatic investigations towards potential translation to early diagnosis and treatment. Using nanoparticle probes engineered with a variety of physical properties (e.g. size, shape, stiffness) and surface modifications^48-50^, their delivery efficacy and trafficking kinetics can be systematically evaluated and compared to drive effective design of nanomedicine. Moreover, those nanoparticles after extravasation can be continuously tracked over extended period to uncover their behaviours and fates, for instance in the brain parenchyma (Supplementary Movie 3), which is vital for determining their selectivity towards specific cell populations and disease types. Additionally, integration with the multiplexing capability, as recently demonstrated for single particle tracking^51^, will enable studies on the complicated interplay among various types of biomolecules, subcellular structures, and biological networks. Given the prevalence of unknowns around the endothelial transport that may be addressed by this technique, we believe it will provide a powerful tool for vascular-related research and drug delivery, paving the way for innovative therapeutic and diagnostic applications for a range of diseases.

## Methods

### UCNP preparation

Core-shell UCNPs of NaYb_0.96_Tm_0.04_F_4_@NaYF_4_ were synthesized via a two-step growth method reported previously^33,35^. The pristine UCNPs capped in oleic acid (OA-UCNPs) were transferred to hydrophilic phase by modifying their surface with phospholipids or polysorbates (PS80). For phospholipid coating, 5 mg OA-UCNPs suspended in 1 mL of chloroform was mixed with 1 mL of chloroform solution containing 12.5 mg of phospholipids (DSPE-PEG_2000_-COOH, Ponsure Biotech) and transferred to a 20 mL glass bottle. The bottle was left open in a fume hood at room temperature for 1 day, and then placed in a vacuum oven at 40°C for 1 hour to completely evaporate the chloroform. The obtained film was hydrated with 5 mL Milli-Q water, followed by vigorous sonication and then filtering through a 0.22 µm PVDF filter to remove any large aggregates. The obtained phospholipid-coated UCNPs were collected by high-speed centrifugation (12,000 rpm, 10 min), washed twice with Milli-Q water, and stored at 4 °C for further use. For PS80 functionalisation, 100 μL of PS80 (Sigma-Aldrich) was added to a 50 mL flask containing 5 mg OA-UCNPs suspended in 2 mL of chloroform. The mixture was stirred for 1 hour at room temperature. Then, 10 mL of Milli-Q water was added to the flask, and the mixture was heated in an oil bath at 70 °C for 4 hours to evaporate the chloroform. During this process, PS80 gradually coated onto the surface of the UCNPs, transferring them into the hydrophilic phase. A 0.22 µm PVDF filter was then used to remove any large aggregates, and the obtained PS80-functionalised UCNPs were stored at 4 °C for further use.

### UCNP characterisation

The nanoparticle size and morphology were measured on a Philips CM10 transmission electron microscope (TEM) operating at 100 kV. Photoluminescence spectroscopy under 980 nm laser excitation was recorded using Fluorolog-Tau3 (Jobin Yvon-Horiba), showing major emission at the 800 nm NIR band. For hydrophilic UCNPs, the nanoparticles were incubated in full DMEM medium supplemented by 10% FBS (Gibco™, Thermo Fisher) for 4 hours, followed by drop-casting onto carbon-coated copper grids (ProSciTech) and drying for 24 h. Concentration of the hydrophilic UCNPs in Milli-Q water was calibrated using NanoSight NS300 (Malvern), and their hydrodynamic size and zeta potential were measured using Zetasizer APS (Malvern). The results of UCNP characterisation are summarised in Supplementary Figures 3 & 4.

### Imaging platform

Supplementary Figure 5 shows the dual-modal fluorescence microscopy platform developed for this study based on a two-deck Olympus IX83 frame. Its upper deck was configured for the confocal mode, and the lower deck for the wide-field mode. To enable simultaneous operation, a dichroic mirror (T750lpxrxt-UF2, Chroma) was inserted in the upper deck to separate visible fluorescence from the zebrafish vasculature and conventional dyes in the confocal mode from NIR upconversion luminescence for tracking UCNPs in the wide-field mode. For the latter, Köhler illumination was configured over the entire field-of-view with a fibre-coupled 980-nm laser (LE-LS-980-500TFCA, LEOPTICS) under an oil-immersion objective (UPLSAPO60XS2, Olympus), while the excitation and emission light were separated by a dichroic mirror (ZT1064rdc-sp, Chroma) and an additional emission filter (FF01-850/SP-25, Semrock). To avoid thermal effects on live zebrafish, the 980-nm laser was pulsed at 10 kHz with 50% duty cycle. Wide-field images were captured using an EMCCD camera (Evolve 512 Delta, Photometrics), while confocal imaging was realised by the Olympus FV3000 laser scanning module using its FluoView control software. When operated simultaneously, the two modalities acquired images on the same focal plane. The Z focus position was indicated by the FluoView software in real time as being adjusted, either in FluoView or directly using the focus knob. Besides, the confocal module was also equipped with a 980-nm laser (VLSS-980-B-900-FA, Connet) for imaging UCNPs when the wide-field mode was not in use and the upper deck was switched to a normal mirror, allowing cross-validation of the 3D distribution of the UCNPs with respect to the vasculature. Laser ablation of the blood vessel can be achieved by line scan in the confocal mode with the in-built 405-nm laser.

### Zebrafish model

A transgenic line [Tg(fli1a:EGFP)] of zebrafish (*Danio rerio*) at 3–5 days post fertilisation was used in the study. The fish was housed in the Zebrafish Facility at Macquarie University, and experiments were conducted under the approval of Macquarie University Animal Ethics (2012/050 and 2015/033). The fish were maintained at 28 °C in a 13 h light and 11 h dark cycle, and embryos were collected by natural spawning and raised at 28.5 °C in E3 medium (5 mM NaCl, 0.17 mM KCl, 0.33 mM CaCl_2_ and 0.33 mM MgSO_4_ buffered to 7.3 pH using carbonate hardness generator (Aquasonic), no methylene blue) according to standard protocols. A solution of 0.003% 1-phenyl-2-thiourea (Sigma-Aldrich) was added around 20 to 24 hours post fertilisation, and the medium was changed every two days to suppress the formation of pigments.

### Dynamic imaging of UCNPs and dyes in live zebrafish

Anesthetised zebrafish were mounted in low-melting agarose (Thermo Fisher) at a concentration of 1% in E3 medium. To visualise the injection process and compare the extravasation between contrast agents, tetramethylrhodamine isothiocyanate labelled dextran (dextran-TRITC, 10 kDa, Thermo Fisher) or Alexa Fluor 647 conjugated bovine serum albumin (BSA-AF647, 66 kDa, Thermo Fisher) at a final concentration of 50 µg/mL were added to the UCNP suspension. Borosilicate glass capillary tube (1.0 mm O.D., 0.78 mm I.D.; Harvard Apparatus) was pulled using a needle puller to generate needles with long, tapered shafts suitable for precise injection. A microinjection apparatus (Picospritzer II, General Valve Corporation) fitted with the pulled glass capillary needle was used to deliver the UCNP suspension (∼5 nL containing around 1–2×10^5^ UCNPs per injection) into the posterior cardinal vein of zebrafish near the sphincter region. The fish were immediately used for visualisation under the purpose-built fluorescence microscope.

To visualise potential extravasation, the fish first underwent rapid surveillance to locate a potential leakage site suggested by any UCNPs outside the vasculature or appearing arrested on the wall of the blood vessel. A confocal Z scan was then performed to record the surrounding environment in 3D, followed by continuous wide-field imaging at the same location while manually keeping the particle of interest in focus, if applicable. Considering the unpredictable nature of nanoparticle extravasation, each location was recorded at video rate for 20 minutes, and the confocal Z scan was repeated afterwards to verify any potential change in the surroundings. The fish was then surveyed again for other potential leakage site. The extravasation of dextran-TRITC and BSA-AF647 was observed during the confocal scans at different time points, as shown in Supplementary Figure 6. The 3D intensity profiles in Fig. 3b–c were obtained using ImageJ.

### Single particle tracking analysis

Image stacks containing potential nanoparticle extravasation events were imported into the TrackIt analysis framework^52^, which extracted the 2D coordinates of the nanoparticles in each frame for subsequent calculation of the key metrics in MATLAB, including displacement over time, instantaneous speed, and mean squared displacement. A moving-average filter was applied to compensate potential rhythmic motion caused by the zebrafish heartbeat. A 6th-order low-pass filter was further applied to the speed data in order to identify phases of movement within the trajectory. Angle wrapping was performed to limit the direction jump between 0 to 180°, allowing segments of directed motion to be clearly distinguished. Sample stage and objective drifts were found minimal during the course of imaging and therefore not accounted for.

### Co-registration of UCNPs and zebrafish vasculature

To facilitate precise co-registration, the field-of-views under the confocal and the wide-field mode were calibrated using a reticle target (R1L3S3P, Thorlabs), as shown in Supplementary Figure 5. In particular, the laser scanning area was carefully adjusted in the confocal software FluoView, so that the two field-of-views matched one another. Using image cross-correlation, the difference in scaling was found to be less than 0.1% and the misalignment less than 2 pixels in XY. The confocal *Z*-stacks were then colour-coded in depth, before being superimposed on the wide-field *t*-series to generate the movies using customised MATLAB codes.

### Evaluation of nanoparticle biodistribution in zebrafish brain

After live imaging, zebrafish injected with UCNPs were released into the E3 medium. At 4 hours post-injection, the fish was anesthetized and fixed by 4% paraformaldehyde (Electron Microscopy Sciences) in phosphate-buffered saline (Thermo Fisher) at room temperature for 30 minutes, followed by rinse in E3 medium and embedding in 1% agarose to examine the 3D distribution of the UCNPs with respect to the vasculature. The data were processed using the Imaris software to calculate the amount of UCNPs outside the blood vessels, as illustrated in Supplementary Figure 7.

## Acknowledgements

The authors acknowledge financial support from the Australian Research Council ARC Discovery Project (DP230102459; DP210103469), NHMRC Leadership Fellowship (B.S.), Cancer Institute NSW Early Career Fellowship (2022/ECF1449), NIMH, NINDS and NIDA support (R21DA056320, M.M.), and Macquarie University Research Fellowship (J.L.).

## Author contributions

G.W. and Y.C. contributed equally to this work. Conceptualisation: Y.L., B.S.; Methodology: Y.L., X.Z., G.W., Y.C.; Investigation: G.W., Y.C.; Formal analysis: Y.C., G.W., J.L., Y.L.; Resources: Y.C., G.W., M.M., P.R.; Visualisation: Y.C., G.W., Y.L.; Supervision: Y.L., B.S.; Writing – original draft: Y.C., G.W., Y.L.; Writing – review & editing: Y.C., G.W., Y.L., J.L., M.M., B.S.

