## Supplementary Information for "Visualising sub-second dynamics of nanoparticle extravasation *in vivo*"

### Captions of Supplementary Movies

Supplementary Movie 1 shows a representative extravasation event in the brain of a 3dpf zebrafish larva, observed continuously for >10 minutes. The upconversion nanoparticles are rendered in magenta, while the blood vessels are colour-coded (indicating the depth) to provide the spatial context of the nanoparticle movement. The time of each frame since the start of the video is annotated at the top. Part of the video has been skipped when the nanoparticle of interest underwent minimum movement. Scale bar: 20 μm.

Supplementary Movie 2 shows a representative extravasation event observed in the truck of a 3 dpf zebrafish larva. The white arrow points to the location where the nanoparticle of interest extravasated, occurring around 3 seconds since the start of the video. Scale bar: 20 μm.

Supplementary Movie 3 shows the movement of upconversion nanoparticles in the brain of a 3 dpf zebrafish larva after extravasation. Scale bar: 10 µm.

Supplementary Movie 4 shows upconversion nanoparticles moving in the *in vitro* BBB model consisting of human brain capillary endothelial cells cultured in a microfluidic chip. The white arrow indicates the location where the nanoparticle of interest exits the apical chamber (in which the cells were cultured) through a slit connecting to the basolateral chamber. Part of the video has been skipped when the nanoparticle of interest underwent minimum movement. Scale bar: 20 µm.

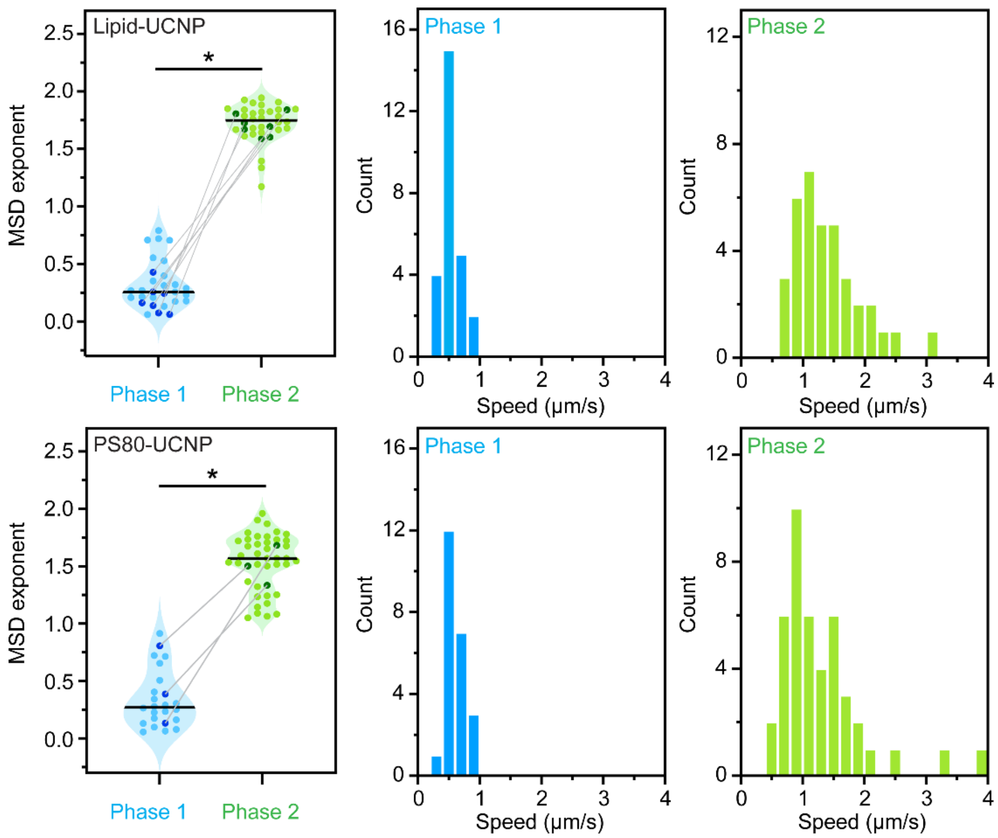


**Supplementary Figure 1 | Statistical analysis of the UCNP trajectories captured in the brain,** including scatter plots of the MSD exponent and histograms of the average speed during the two phases, for both phospholipid-coated and PS80-functionalised particles. The pairs of darkened data points connected by grey lines in the scatter plots represent full extravasation events of which the entire process was observed, while others are calculated from recorded nanoparticle trajectories moving either along or outside the vessels to supplement the limited number of full events. The asterisks represent $p < 0.05$ for Mann-Whitney test to indicate statistical difference.

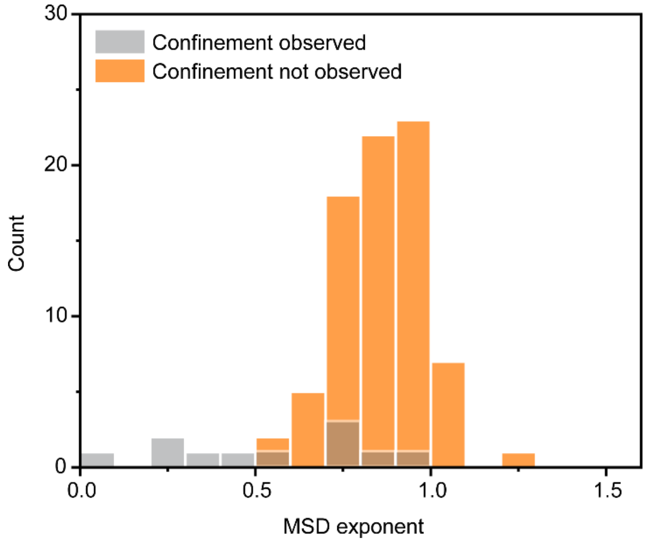


**Supplementary Figure 2 | Histograms of the MSD exponent obtained for UCNPs moving outside the intersegmental vessels in the trunk,** with the majority appearing in free diffusion (orange bars) while some exhibiting confined motion (grey bars). The latter, as well as the mean MSD exponent being slightly below 1 for the former, suggests limited interstitial space outside the vasculature.

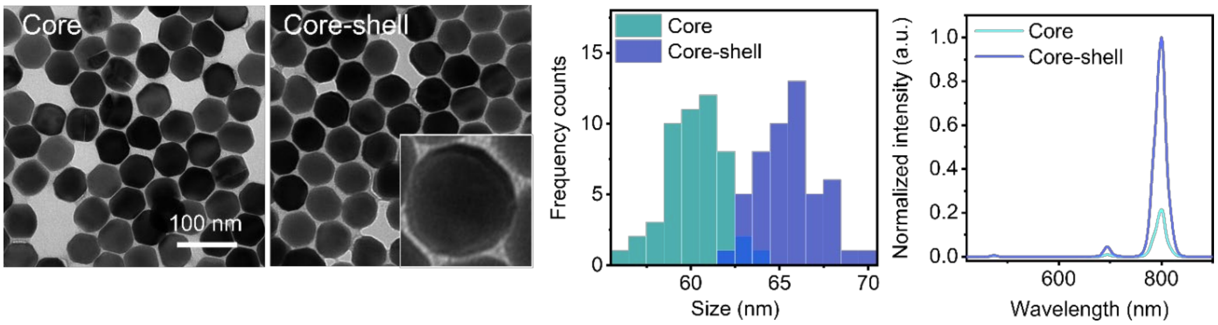


**Supplementary Figure 3 | Characterisation of OA-UCNPs,** including TEM of core and core-shell nanoparticles, size distributions, and emission spectra under 980 nm laser excitation at 10 $W/cm^2$.

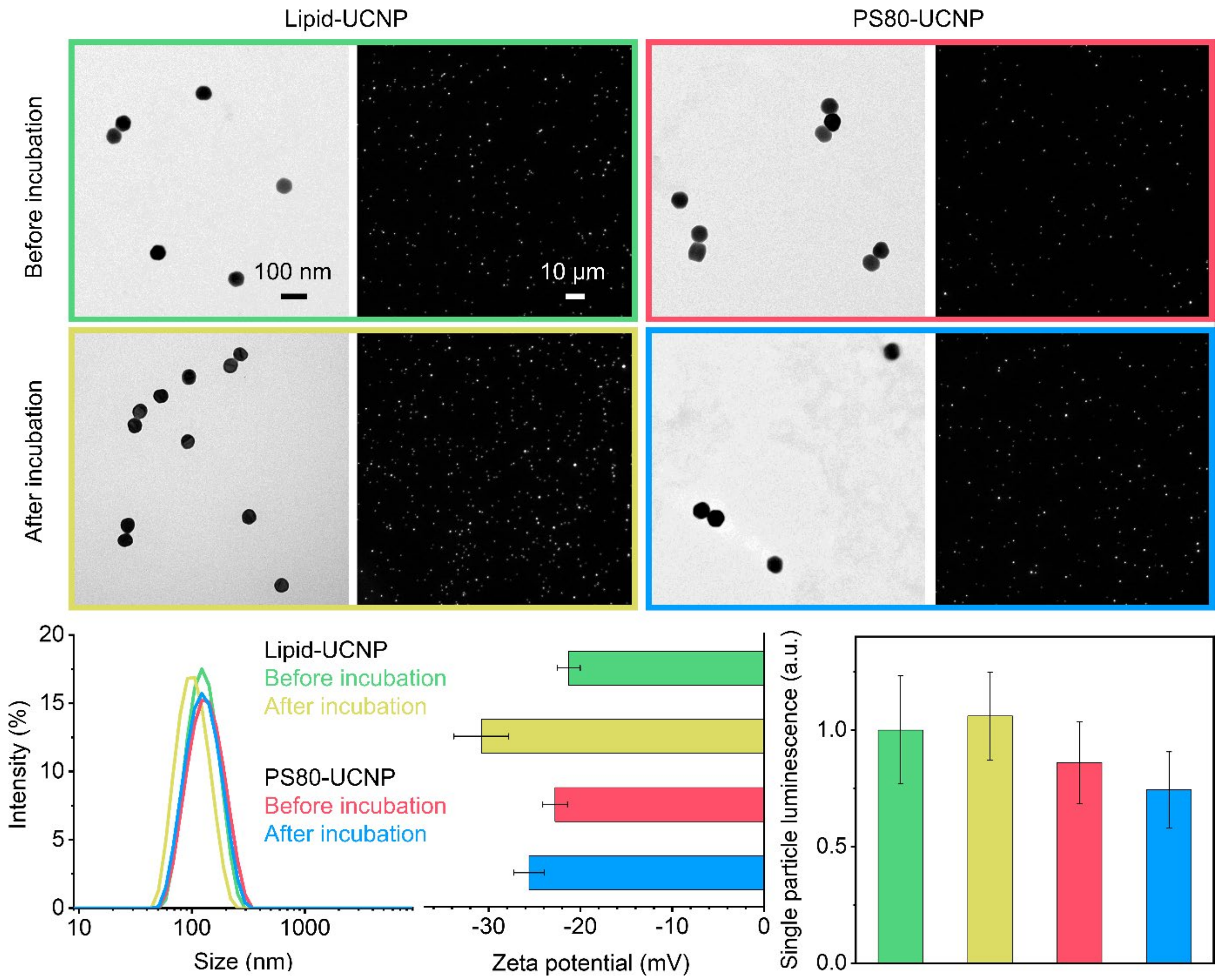


**Supplementary Figure 4 | Characterisation of hydrophilic UCNPs before and after incubation with biological fluids,** including TEM, wide-field luminescence images under 980 nm excitation, hydrodynamic diameters, zeta potential, and relative luminescence intensities measured from the dim objects assumed to be individual UCNPs. Error bars represent standard deviation.

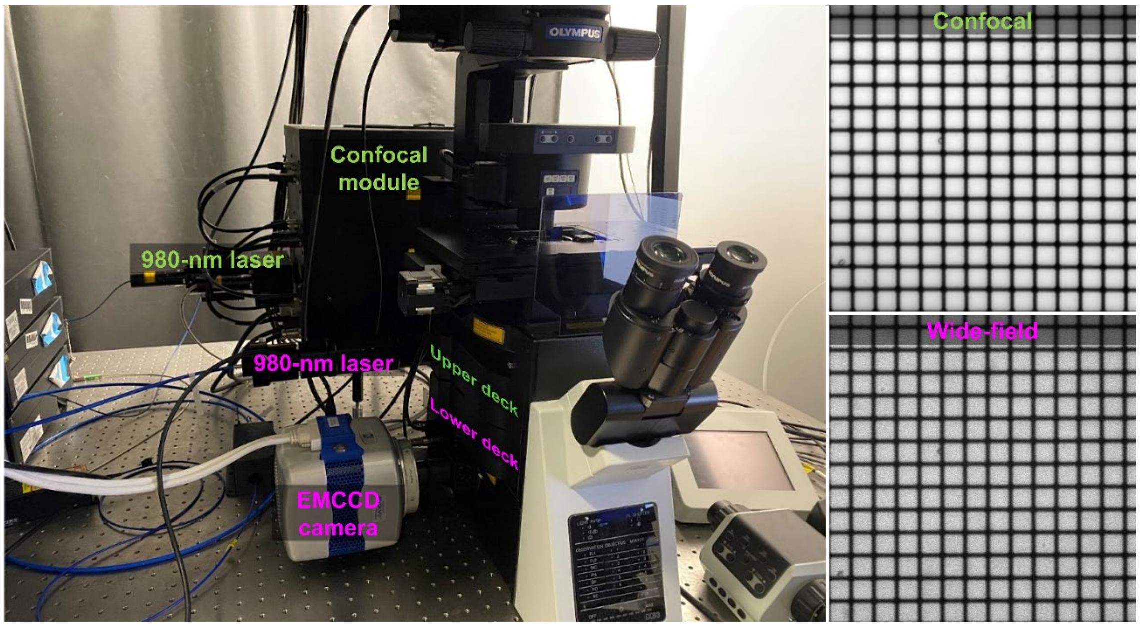


**Supplementary Figure 5 | Dual-modal fluorescence microscopy platform.** Key parts for the confocal and the wide-field mode are marked in lime and pink colours in the photo. The field-of-views captured in each mode on a reticle target with 10 µm intervals under a 60X objective are presented on the right.

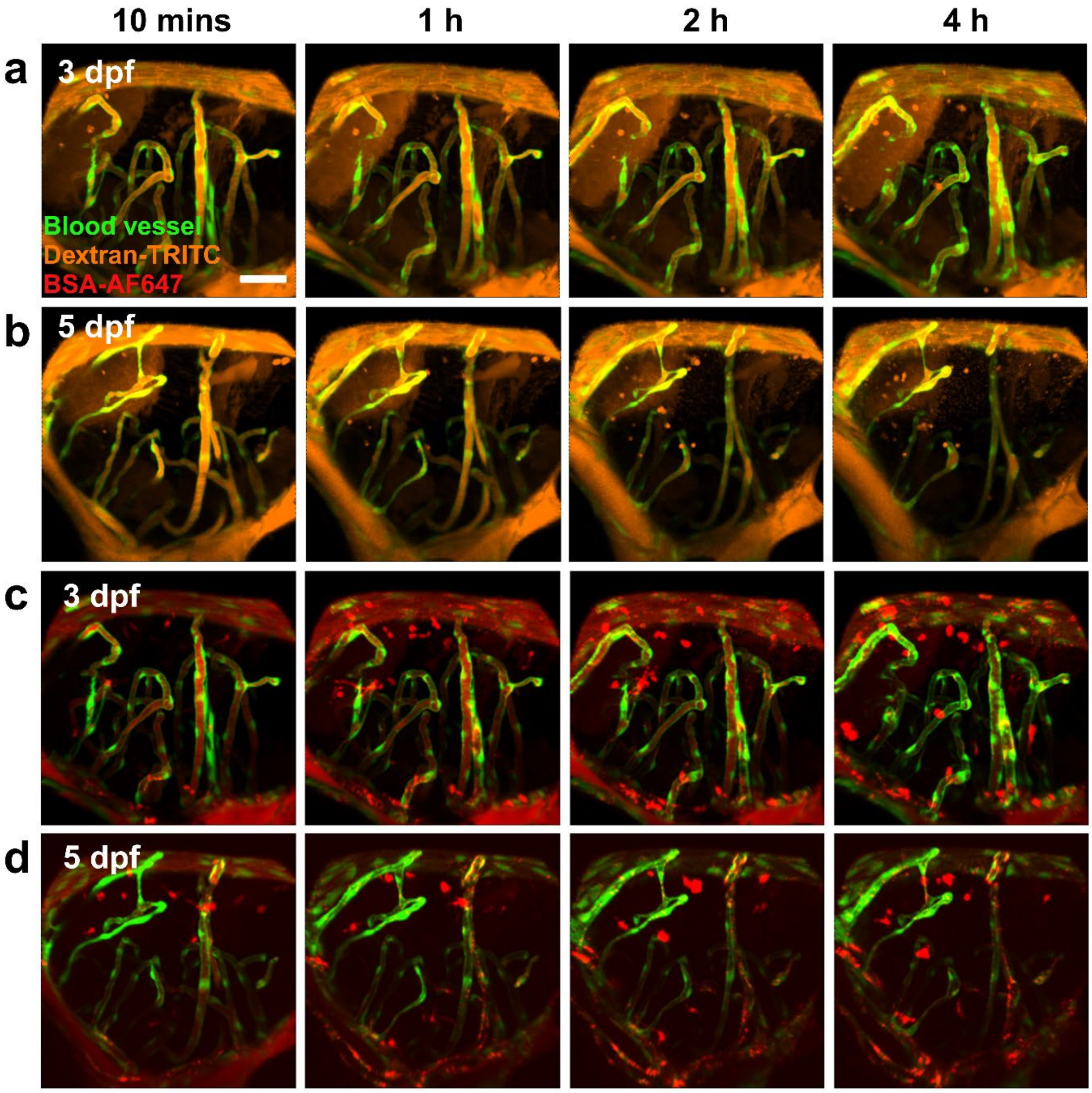


**Supplementary Figure 6 | Extravasation of fluorescent probes in the brain of zebrafish larvae at different time points.** (a, b) Representative confocal images of dextran-TRITC distributed in the brain of 3 dpf and 5 dpf zebrafish, respectively. (c, d) Representative confocal images of BSA-AF647 distributed in the brain of 3 dpf and 5 dpf zebrafish, respectively. Scale bar: 50 µm.

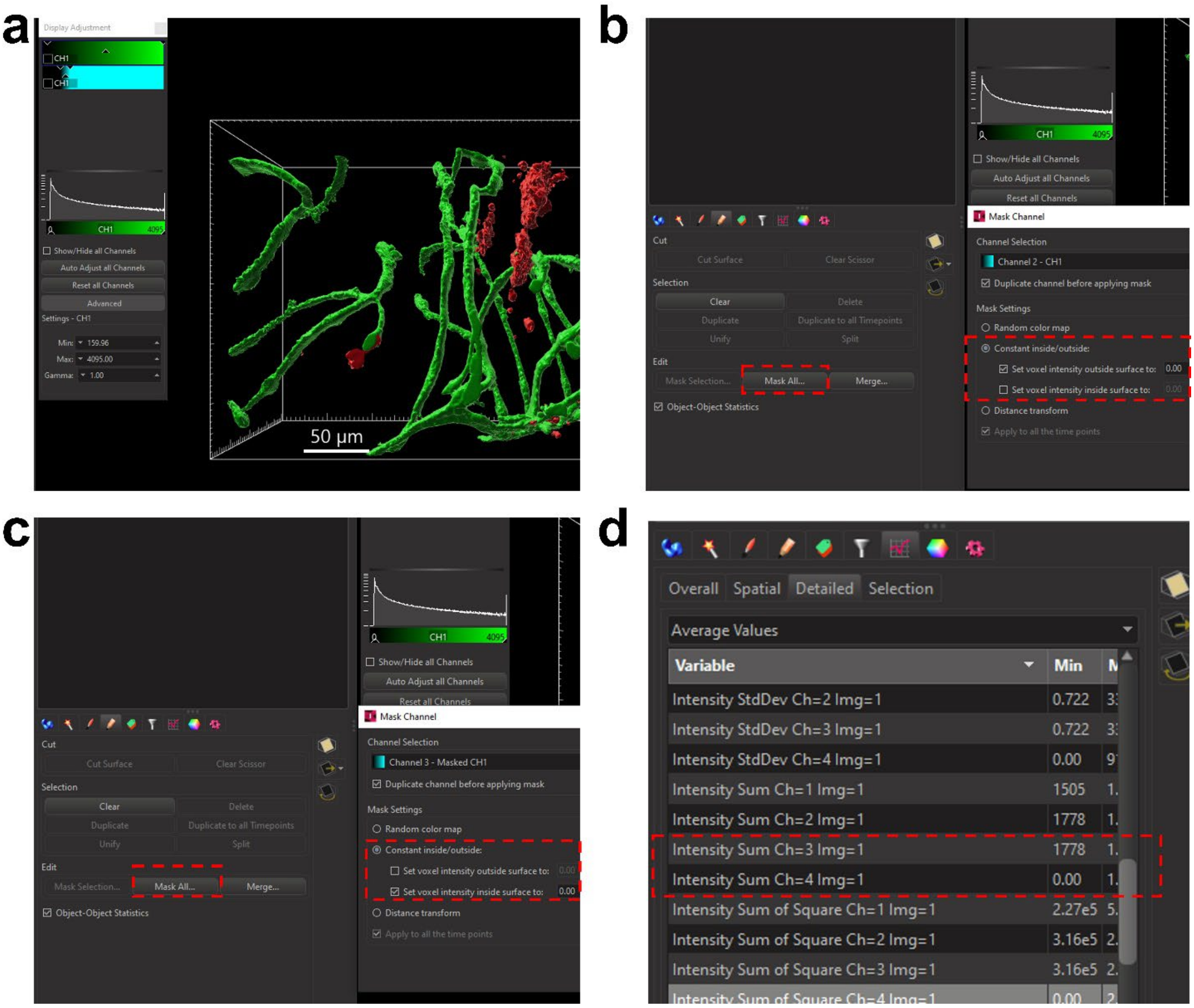


**Supplementary Figure 7 | Procedures to evaluate UCNP distribution in zebrafish brain region.** (a) 3D rendering of the Z-stack images, with the green channel indicating blood vessels and the red channel for UCNPs. (b) The option 'Mask All' is applied to select the voxels corresponding to the UCNPs (as Channel 3). (c) The option 'Mask All' is applied to select the voxels correlated to and inside the blood vessels, so that these voxels are discarded for subsequent calculation of the UCNP signal outside the vessel (as Channel 4). (d) Intensity is then summated over Channel 3 to represent the total UCNPs in the brain region, and over Channel 4 for the UCNPs outside the brain vasculature.

**Supplementary Note 1. Imaging performance for dynamic tracking of single UCNPs**

A glass coverslip containing monodispersed UCNPs was prepared to verify the single nanoparticle imaging capability. Due to optical diffraction, one UCNP was projected to multiple pixels (Supplementary Fig. 8a), which can be described by the point spread function (PSF) assumed as Gaussian,

$$\mathrm{PSF}(x) = ae^{-\frac{(x-\mu)^2}{2w^2}} + b \tag{S1}$$

Parameters of the Gaussian function $a$, $b$, $\mu$ and $w$ can be fitted from the experimental data, as shown in Supplementary Fig. 8b. In particular, $w = 1.3$ (pixel), indicating the size of the PSF. Accordingly, for each particle, we select an area of 5×5 pixels (± 1.9$w$, ~94%) and subtract the pixel bias ($b = 400$) to calculate its integrated intensity. This was conducted for 100 single UCNPs (Supplementary Fig. 8c), resulting in a mean integrated intensity of $1.4\times10^4$ with standard deviation of $3.3\times10^3$ (Supplementary Fig. 8d).

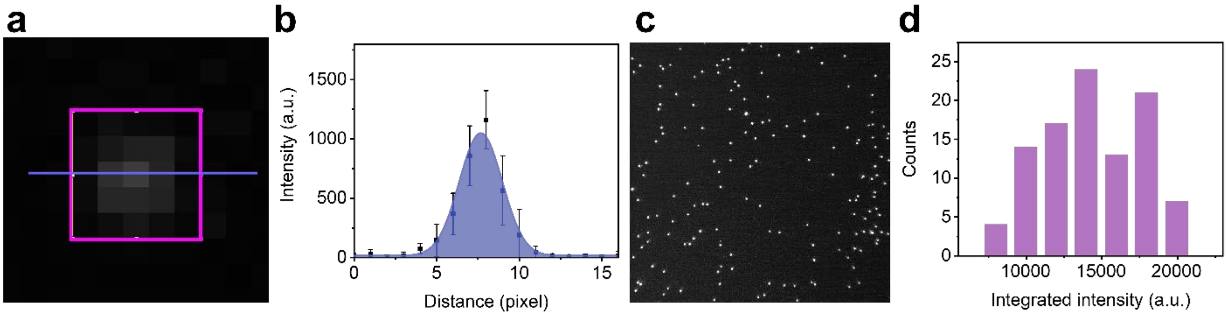


**Supplementary Figure 8 | Luminescence intensity from single UCNPs.** (a) Pixels corresponding to a single particle. (b) Gaussian fitting of the signal profile along the line in (a). (c) A field-of-view of the mono-dispersed UCNPs under wide-field imaging. (d) Intensity histogram of single UCNPs.

The experimental result above was then validated against theoretical analysis. For a single UCNP of diameter $d$ and unit cell volume $V_U$, the number of lanthanide ions $N_{Ln}$ is given by

$$N_{\mathrm{Ln}} = Z\frac{\pi d^3}{6V_{\mathrm{U}}} \tag{S2}$$

in which $Z = 1.5$ indicates on average 1.5 lanthanide ions per unit cell. For the UCNPs used here, $d =$ 60 nm. Since the Tm doping concentration (4%) is small, the unit cell volume can be approximated by that of $NaYbF_4$, $V_U = 0.106$ nm$^3$. Therefore, we estimate the numbers of $Yb^{3+}$ sensitizers $N_S =$ $1.53\times10^6$ as well as of $Tm^{3+}$ activators $N_A = 6.4\times10^4$ in a single nanoparticle.

The photon absorption rate of a single UCNP can be expressed as

$$q_{\mathrm{ab}} = N_{\mathrm{s}}\sigma\frac{I_{\mathrm{ex}}\lambda_{\mathrm{ex}}}{hc} \tag{S3}$$

in which $\sigma = 1\times10^{-20}$ cm$^2$ is the absorption cross-section of the $Yb^{3+}$ sensitizers, $I_{ex}$ is the excitation irradiance, $\lambda_{ex} = 980$ nm is the excitation wavelength, $h$ is the Planck constant, and $c$ is the speed of light. Denoting the quantum yield of the upconversion emission as $\gamma$, we then obtained the emission rate of a single UCNP as

$$q_{\text{em}} = \gamma q_{\text{ab}} \tag{S4}$$

In our experiment, the excitation source provides continuous-wave power of 0.6 W, but has been operated under pulsed mode with 50% duty cycle. The field-of-view under the microscope objective is $1.05\times10^{-3}$ cm$^2$. Thus, $I_{ex} = 286$ W cm$^{-2}$, and consequently $q_{ab} = 2.16\times10^7$ (photon) s$^{-1}$. Assuming $\gamma = 0.1\%$ for the 800 nm emission band, we estimate $q_{em} = 2.16\times10^4$ (photon) s$^{-1}$.

To produce a luminescence image, the photons emitted by a single UCNP are projected onto the pixels of an image sensor (camera), generating photoelectrons that are then digitised to an intensity value for each pixel. Assuming light collection efficiency $\alpha$ and quantum efficiency of the camera $\beta$, the photoelectron rate (integrated over all pixels corresponding to a single UCNP) is expressed as

$$q_{\text{s}} = \alpha\beta q_{\text{em}} \tag{S5}$$

For our imaging system, we estimate $\alpha = 0.2$ based on numerical aperture of 1.3 and transmittance of 0.75 at 800 nm for the objective lens as well as transmittance of 0.9 for other optical elements. The system is coupled to an electron-multiplying charge-coupled device (EMCCD) camera, with $\beta = 0.85$ according to the specification. Thus, $q_s = 3.67\times10^3$ (e$^-$) s$^{-1}$. The EMCCD further amplifies the photoelectrons in each pixel well before converting to intensity,

$$A = q_{\text{s}} T G \Phi \tag{S6}$$

in which $T$ is the exposure time, $G$ is the electron-multiplying (EM) gain, and $\Phi$ is the digitisation factor expressed as

$$\Phi = \frac{1}{C}(2^M - 1) \tag{S7}$$

where $C$ is the EM register full-well, and $M$ is the bit depth of the analogue-to-digital converter. In our experiment, $G = 1000$, $C = 8\times10^5$ (e$^-$), and $M = 16$. Thus, with exposure $T = 50$ ms, we obtain $A = 1.5\times10^4$ for a single UCNP, which is in excellent agreement with the experimental result.

Apart from the signal, detection of single UCNPs is also influenced by the noise, including optical noise and electronic noise. The former is negligible since the anti-Stokes emission from UCNPs can be easily distinguished from biological autofluorescence and residual scattering. The latter of each pixel, represented by variance $\sigma_{\rm pix}^2$, consists of three contributions from the signal $\sigma_{\rm s}^2$, from the dark current $\sigma_{\rm d}^2$, and from the readout noise $\sigma_{\rm r}^2$,

$$\sigma_{\rm pix}{}^2 = \sigma_{\rm s}^2 + \sigma_{\rm d}^2 + \sigma_{\rm r}^2 \tag{S8}$$

Assuming the signal and the dark current follow Poisson statistics, we then have

$$\sigma_{\rm pix}{}^2 = q_{\rm pix}TG^2F^2\Phi^2 + q_{\rm d}TG^2F^2\Phi^2 + n_{\rm r}^2\Phi^2 \tag{S9}$$

in which $q_{\rm pix}$ is the photoelectron rate (signal) and $q_{\rm d}$ is the dark current of the pixel (in $e^-$/s), and $n_r$ is the read noise (in $e^-$ rms). $F$ is the noise factor introduced by the EM gain process, often assumed $F^2 = 2$ for an EMCCD camera. Thus, the peak signal-to-noise ratio (SNR) can be calculated as

$$\mathrm{SNR} = \frac{A_{\rm pix}}{\sigma_{\rm pix}} = \frac{q_{\rm pix}TG\Phi}{\sqrt{q_{\rm pix}TG^2F^2\Phi^2 + q_{\rm d}TG^2F^2\Phi^2 + n_{\rm r}^2\Phi^2}} = \frac{q_{\rm pix}T}{\sqrt{(q_{\rm pix} + q_{\rm d})TF^2 + \frac{n_{\rm r}^2}{G^2}}} \tag{S10}$$

in which $A_{\rm pix}$ is the peak pixel intensity from a single UCNP. As we use the PSF in Eq. (S1) to describe the image of a single UCNP, $q_{\rm pix}$ is a fraction of $q_s$,

$$q_{\rm pix} = \xi q_s \tag{S11}$$

For the pixel in the middle (i.e. ± 0.5 pixel), the fraction can be calculated as $\xi = 0.3$, corresponding to ± 0.38$w$ (since $w$ = 1.3 pixel). Hence, $q_{\rm pix} = 1.1\times10^3$ $e^-$/s for the middle pixel. By contrast, according to the specification of the EMCCD camera, $q_{\rm d}$ = 0.003 $e^-$/pixel/s, and $n_{\rm r} < 1$ $e^-$ with EM gain enabled. Therefore, both terms can be ignored, and we obtain

$$\mathrm{SNR} = \sqrt{\frac{\xi q_s T}{F^2}} \tag{S12}$$

By substituting the symbols with their values obtained above, SNR = 5.25, which satisfied the Rose criterion for identification of individual nanoparticles with certainty.

It should be noted that, at given exposure time, SNR is primarily determined by the brightness of the nanoparticles. For UCNPs, the single-particle brightness is influenced by both the nanoparticle size (presumably proportional to the cubic) and the excitation irradiance (presumably proportional to the square for the two-photon process behind the main emission band at 800 nm). Therefore, it is possible to use smaller UCNPs as long as adequate SNR can be maintained by increasing the excitation irradiance.

We also validated the single particle tracking capability by imaging the diffusion of phospholipid-coated UCNPs in the medium. For Brownian motion in two dimensions, the diffusion coefficient $D$ of a particle can be determined from the Mean Squared Displacement (MSD) of the trajectory, given by

$$\mathrm{MSD}(\Delta t) = 4D\Delta t \tag{S13}$$

in which $\Delta t$ is the time interval of the frames. Based on the experimental data, we obtained $D = 2.45 \pm 0.76$ μm$^2$/s.

In comparison, the diffusion coefficient $D$ can also be calculated theoretically using the Stokes–Einstein–Sutherland equation,

$$D = \frac{k_B \Theta}{6\pi\eta r} \tag{S14}$$

where $k_B$ is the Boltzmann constant ($1.38\times10^{-23}$ J/K), $\Theta$ is the absolute temperature, $\eta$ is the viscosity of the medium, and $r$ is hydrodynamic radius of the nanoparticles. Using $\Theta = 300$ K, $\eta = 1.5$ cP, and $r = 60$ nm (cf. Supplementary Fig. 1e), we obtained $D = 2.44$ μm$^2$/s, offering excellent agreement to the experimental result above.

**Supplementary Note 2. Visualising UCNPs in a microfluidic BBB model**

For proof-of-concept, the new technique was applied to an *in vitro* BBB model first prior to the *in vivo* experiments in living zebrafish. Human brain capillary endothelial cells (hCMEC/D3, CLU512, Cedarlane) were maintained in endothelial cell basal medium (EMB-2, Lonza), supplemented with EGMTM-2 Endothelial SingleQuots$^{TM}$ Kit (Lonza), and grown in flasks coated with gelatine (10 mg/mL, Sigma). A microfluidic chip (SynBBB IMN2 radial, SynVivo) was used to build the *in vitro* model, with its apical channels coated with human fibronectin (50 µg/mL, Sigma) and incubated at 37°C for 1 hour prior to cell seeding. The hCMEC/D3 cells at a concentration of 3–5×10$^{7}$ cells/mL in EMB-2 medium were seeded into the fibronectin-coated apical channels at a flow rate of 10 µL/min using a syringe pump. The chip was then inverted and incubated for 4 hours to allow cells to adhere to and grow on the top surface of the channels. After that, the same cell seeding procedure was repeated, with the chip placed in the normal orientation to enable cell growth on the bottom surface of the channels. The chip was then left in the incubator overnight. On the next day, EMB-2 media flow (0.1 µL /min) was applied to the apical channels using a syringe pump (Fusion 200, Chemyx) while the setup was covered with aluminium foil to avoid light exposure. After another 24 hours of culture, the chip was ready for the imaging experiment.

To verify the successful development of the *in vitro* model, the cells inside the chip were fixed in 4% paraformaldehyde (in PBS) at room temperature for 10 minutes, and then permeabilized with 0.1% Triton X-100 (in Tris-buffered saline, TBS) for 10 minutes followed by blocking with 3% bovine serum albumin in 1× TBS plus Tween-20 (TBST) for 1 hour at room temperature. Afterwards, primary antibody for the endothelial tight junctions (anti-VE-Cadherin, 1:200 dilution, D87F2, Cell Signaling Technology) in the above blocking buffer was applied, and the chip was incubated overnight at 4°C. The chip was washed twice with TBST and twice with TBS (5 minutes each wash), then incubated with secondary antibody (Alexa Fluor 488-conjugated Goat Anti-Rabbit IgG, 1:200 dilution, Thermo Fisher) for 2 hours at room temperature in the dark. Following secondary antibody incubation, the chip was again washed twice with TBST and twice with TBS (5 minutes each). Cell nuclei was stained with Hoechst 33342 (1:2000 dilution in PBS, Thermo Fisher) and cell membranes were counterstained using CellMask™ Plasma Membrane Stain (in PBS, C10045, Thermo Fisher) for 10 minutes at room temperature. Imaging was performed using a Zeiss 880 confocal microscope (Supplementary Fig. 9a-c).

After the protocol was confirmed, freshly prepared chips were mounted on the purpose-built microscopy platform, which was equipped with a temperature-controlled plate, $CO_2$ supply, and humidity regulation to maintain the cell culture condition. Dextran-TRITC and phospholipid-coated

UCNPs were dispersed in the EMB-2 medium with final concentration of 50 µg/mL for the fluorescent probe and approximately $1\times10^8$ particles per mL for the UCNPs. Using a syringe pump, the suspension flew through the microfluidic chip, initially at a flow rate of 0.5 µL/min for 20 minutes to ensure the entire apical channels were filled with the dye and the UCNPs. Afterwards, the flow rate was reduced to 0.1 µL/min, and the UCNPs interacting with the synthetic BBB were continuously imaged at video rate in real time, as shown in Supplementary Movie 4. Besides, fluorescence from dextran-TRITC was collected at selected time points in order to monitor the BBB integrity during the experimental period (Supplementary Fig. 9d).

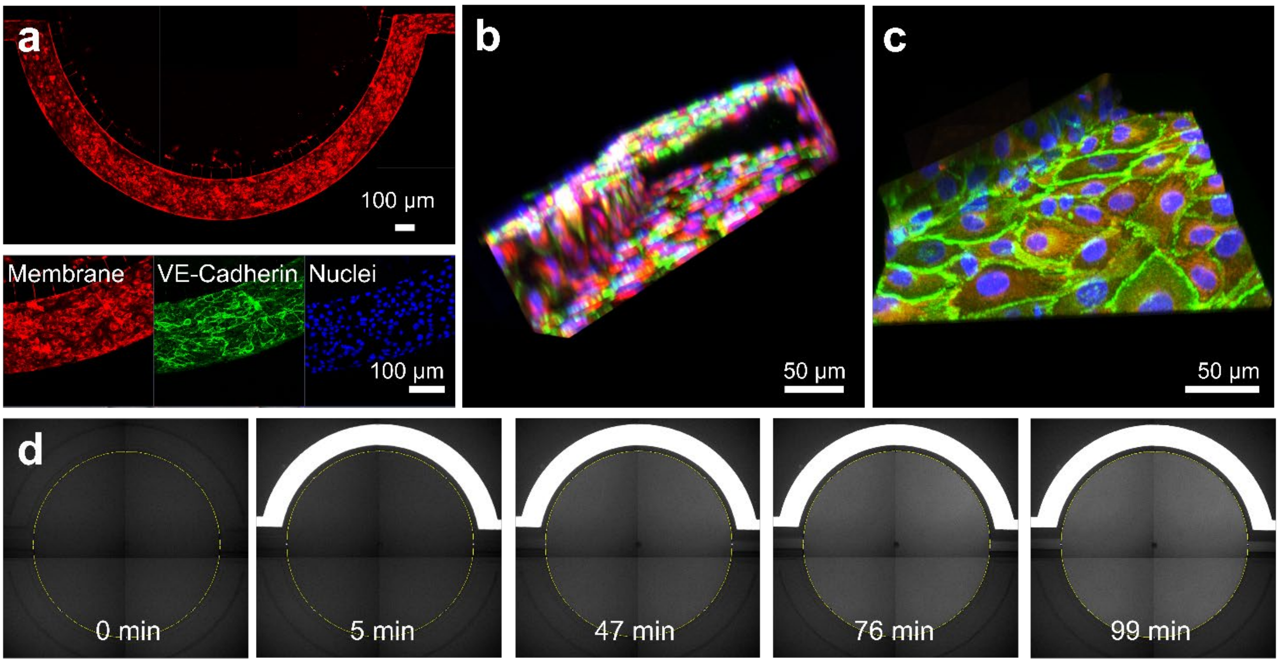


**Supplementary Figure 9 | Validation of the *in vitro* BBB model.** (a) Confocal images of the cell-seeded microfluidic chip, showing cell membrane in red, endothelial tight junctions (labelled with VE-cadherin) in green, and cell nuclei in blue. (b, c) 3D reconstruction of the BBB structure in the apical channel. (d) Evaluation of the BBB integrity by monitoring the diffusion of the dextran-TRITC from the apical channels to the basolateral chamber.
